# Harnessing protein folding neural networks for peptide-protein docking

**DOI:** 10.1101/2021.08.01.454656

**Authors:** Tomer Tsaban, Julia Varga, Orly Avraham, Ziv Ben-Aharon, Alisa Khramushin, Ora Schueler-Furman

## Abstract

Highly accurate protein structure predictions by the recently published deep neural networks such as AlphaFold2 and RoseTTAFold are truly impressive achievements, and will have a tremendous impact far beyond structural biology. If peptide-protein binding can be seen as a final complementing step in the folding of a protein monomer, we reasoned that these approaches might be applicable to the modeling of such interactions. We present a simple implementation of AlphaFold2 to model the structure of peptide-protein interactions, enabled by linking the peptide sequence to the protein c-terminus via a poly glycine linker. We show on a large non-redundant set of 162 peptide-protein complexes that peptide-protein interactions can indeed be modeled accurately. Importantly, prediction is fast and works without multiple sequence alignment information for the peptide partner. We compare performance on a smaller, representative set to the state-of-the-art peptide docking protocol PIPER-FlexPepDock, and describe in detail specific examples that highlight advantages of the two approaches, pointing to possible further improvements and insights in the modeling of peptide-protein interactions. Peptide-mediated interactions play important regulatory roles in functional cells. Thus the present advance holds much promise for significant impact, by bringing into reach a wide range of peptide-protein complexes, and providing important starting points for detailed study and manipulation of many specific interactions.

## Introduction

Peptide-protein interactions are highly abundant in living cells and are important for many biological processes (1). It is estimated that up to 40% of interactions in cells are mediated by peptide-protein interactions, or peptide-like interaction (2): short segments, isolated or embedded within unstructured regions that mediate binding to a partner (3). In addition, peptides are often used for biotechnological applications, drug delivery, imaging, as therapeutic agents and other applications (4–7), by binding proteins and mediating or blocking interactions.

Determining the 3-dimensional structure of these peptide-protein complexes is an important step for their further study. They can provide the basis to identify hotspot residues that are crucial for an interaction (8–10), and mutation of these can be used to study the functional importance of that interaction (11). They also serve as a starting point for the design of strong and stable peptidomimetics (12,13).

However, peptide-mediated interactions pose significant challenges, both for their experimental as well as their computational characterization: On the experimental side, these interactions are in many cases weak, transient, and considerably influenced by their context, resulting in often noisy experiments. For many, the structure cannot be determined by e.g. crystallography. On the computational side, no structure is usually known for the isolated peptide prior to its binding, in contrast to classical domain-domain docking. This significantly complicates peptide-protein docking, in particular blind docking where only the peptide sequence and the receptor structure are known (14). In order to succeed in the study and design of peptide-protein interactions and peptide-protein docking, we must gain a better understanding of the peptide conformational preferences.

Can we learn from existing structures on peptide conformations? Albeit mediating ~40% of the cellular interactions, peptide-protein complexes only compose a small fraction of solved structures. This is due to the small size of the interfaces and the weakness of the interactions. Previous work has demonstrated that peptide-protein interactions are a subset problem of protein folding and form structural interface motifs also found within monomers (15–18). Thus information regarding peptide docking can be extracted from monomeric structures (19,20). Given the scarcity of structural data regarding peptide-protein complexes, incorporating information and knowledge from monomers to understand peptide-protein interactions would be highly impactful.

It was shown that a peptide conformation very similar to the bound conformation is included within a set of sequence-and (predicted) secondary structure-similar fragments extracted from monomer structures (21). Based on this finding, we developed the first high resolution, global blind peptide docking protocol, PIPER-FlexPepDock (PFPD) (21), in which a fragment conformer ensemble extracted from monomer structures (using the Rosetta Fragment Picker, (22)) is used to represent the peptide conformation, exhaustively rigid-body docked onto the receptor (using PIPER FFT-based docking; (21,23)), and finally further refined to atomic resolution with local sampling of both internal peptide and rigid body degrees of freedom (using Rosetta FlexPepDock; (24)).

Beyond the successful representation of peptide conformations by monomer-derived fragment libraries, additional notions regarding peptide folding and peptide-protein interactions can be made. It was shown early that functional proteins can be reconstituted from short fragments of the original sequence, indicating that covalent linkage is not necessarily a prerequisite for monomer folding (16,17). It was also suggested that peptide-protein binding can be seen as a step in monomer folding and complementation (18). Indeed, we and others have been able to model protein-peptide interactions with a protocol that is based on complementation of surface patches, using structural patches derived from interfaces but also from within monomers (19,20). This concept lays the grounds to novel approaches in peptide-protein docking, where information from monomers is efficiently integrated in the search space for peptide docking.

Motivated by initial proofs of the feasibility of global peptide-protein docking with PFPD, numerous computational methods have been developed, with good performance overall (14,25). As with PFPD, many focus on efficient low-resolution docking (26,27), to be combined followed by high resolution refinement (e.g. by Rosetta FlexPepDock). Others leverage information about protein interfaces to find matches for similar interface patches (20,28,29). However, while some interactions are well modeled, there is still much room for improvement of a truly general protocol, as many targets are complicated to dock in the correct manner, in particular if binding is accompanied by large conformational adaptations on the receptor side.

Recent years have seen dramatic acceleration and advances in the field of protein structure prediction (30), as particularly showcased by the last CASP14 experiment (31). The application and development of deep neural network (NN) architectures to predict monomeric protein structures has set the grounds for highly accurate computational models. AlphaFold2 (AF2) developed by Google Deepmind was able to generate models of exceptional resolution approaching for the first time the resolution of crystallography experiments (32). Significantly improved modeling was also reported for RoseTTAFold, the most recent version developed by RosettaCommons, that followed ideas from AF and also implemented fully continuous crosstalk between 1D, 2D and 3D information (33). These advances were possible thanks to (1) the immense increase of sequenced proteins, allowing for the generation of highly informative Multiple Sequence Alignments (MSA), (2) the large amount of solved structures in the Protein Data Bank (PDB) that has been accumulated by low throughput and extensive experimental work for the past decades, and (3) the development of novel NN architectures that provide improved presentation and learning of 3-dimensional features, alongside optimized, parallel computation hardware architectures (30). Importantly and luckily, RoseTTAFold (33) as well as AF2 (32) are now freely available to the scientific community as convenient Google Colab notebooks (34). Moreover Deepmind and the EBI have released a database of AF2 structural models for full proteomes (35). All these provide exciting avenues for new ideas of protocol development, and applications to many biological systems that were not amenable to structural characterization in the past. These are truly exciting times!

Can these NNs also model peptide-protein interactions, and not only monomers? We note that both RoseTTAFold and AF2 NNs were trained on single chain protein structural data, and both use Multiple Sequence Alignments (MSA) as a critical step in structure prediction. Prediction of protein-protein complexes was shown to be possible given an informative MSA (36,37), and it has also been debated whether it is indeed necessary to provide paired sequences for successful extraction of interface information (38). However, creating an effective MSA for the peptide side of a peptide-protein interaction is more challenging.

If peptide-protein interfaces are indeed abundant in monomer structures, and if indeed peptide-protein interactions are part of protein folding as stated above, RoseTTAFold and AF2 should in principle allow also for the modeling of peptide-protein complex structures. Moreover, they could alleviate the lack of data impairing the ability to fully employ DL for peptide-protein interactions.

Here we present a novel method for global peptide-protein docking, which incorporates the biological concept of peptide-protein interactions mimicking protein folding, harnessing neural networks trained to predict protein structure. We assess the protocol on an extensive, non-redundant set of peptide-protein complexes extracted from our AutoPeptiDB database (39) consisting of 162 interactions, each involving a distinct, single ECOD domain. We demonstrate that no multiple sequence alignment of the peptide partner is necessary for this to work. We also provide a detailed comparison of this approach to the currently top performing protocol PFPD (21,29). This is demonstrated on a small, representative, previously well-studied set of protein-peptide interactions, consisting of peptides with and without known binding motifs. Finally, we explore specific types of interactions of special interest, including examples in which peptide binding induces a large conformational shift upon binding. The latter are very challenging to model using docking, but easily amenable to AF2 modeling as a whole.

## Results

### Adaptation of NN-based structure prediction to peptide docking

By adding the peptide sequence via a poly-glycine linker to the C-terminus of the receptor monomer sequence, we mimicked peptide docking as monomer folding (see Methods). A similar approach has been suggested by others (38,40)). This is based on the assumption that the NN should identify the polyglycine segment as non-relevant and use it merely as a connector (**Figure 1E**). We note that in contrast to AF2, a similar tactic using RoseTTAFold did not succeed, but rather attempted to fold the polyglycine into a globular structure or create various loops with intra-loop interactions (**Supplementary Figure S1)**. We therefore proceeded with AF2 for NN-based peptide docking.

**Figure 1.**
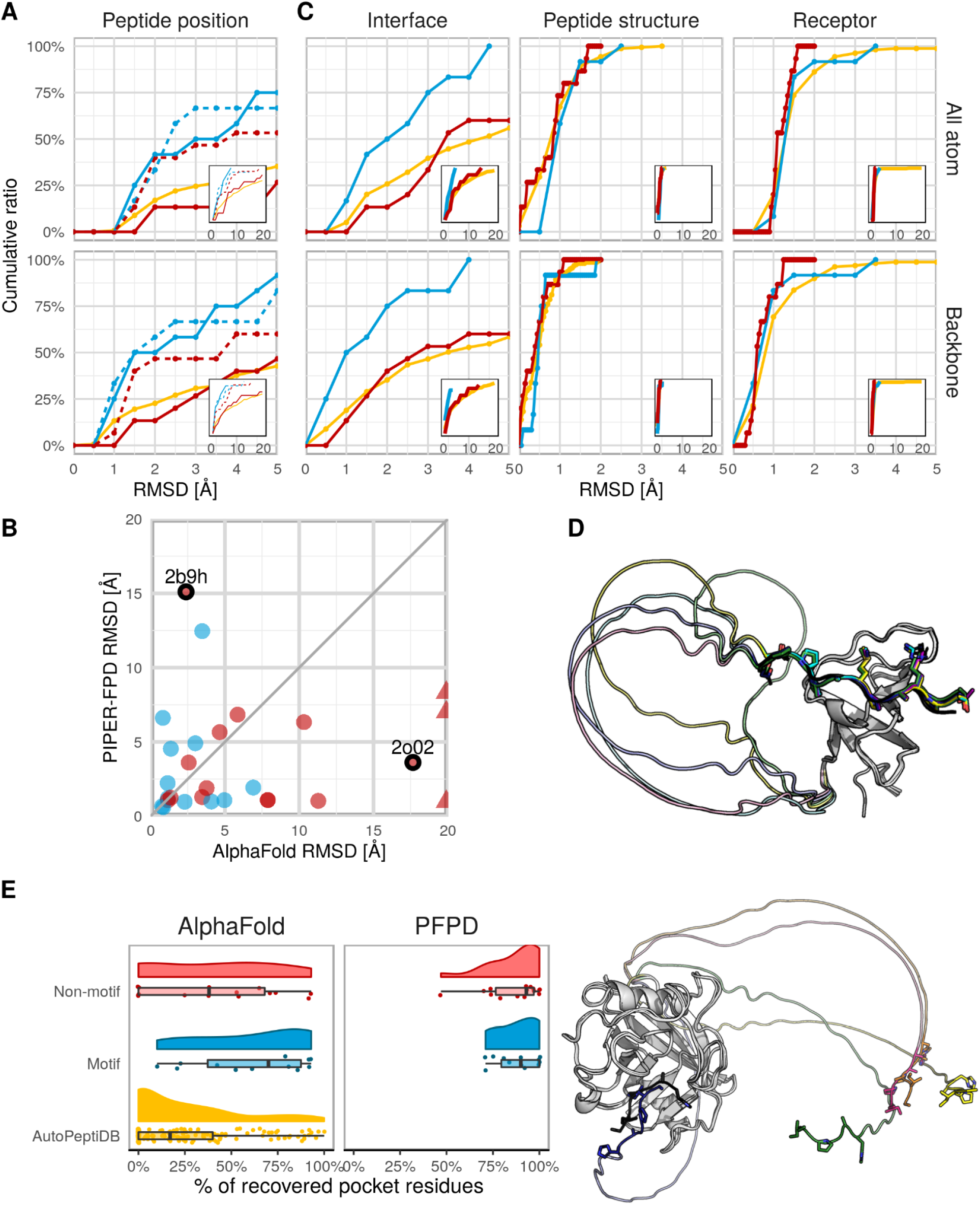
AF2 can be adapted to model peptide-protein interactions at remarkable accuracy. **(A)** Performance of AF2 and PFPD on *motif* and *non-motif PFPD sets*, as measured by the accuracy of the modeled peptide in the binding site (Peptide position; calculated over the interface residues). Cumulative performance plots are shown for RMSD measures over all/backbone peptide interface residue atoms (top/bottom panels, respectively). Performance for AF2 and PFPD is shown in continuous and dashed lines respectively. Results are plotted for the *motif* and *non-motif PFPD sets* in blue and red, respectively (total of 27 complexes). In addition, performance of AF2 is shown for the larger *non-redundant AutoPeptiDB set* in yellow (162 complexes). See **Supplementary Figure S3** for corresponding plots of the distributions of RMSD values. **(B)** Correlation between performance of AF2 and PFPD for the *motif* and *non-motif PFPD sets*. Triangles indicate capped values (>20.0Å RMSD). Complexes discussed further in the text are highlighted with black circles. **(C)** Overall assessment of performance of AF2 for the interface as well as for the individual partners, after each is superimposed separately. **(D)** Examples of successful and failed predictions. For the successful prediction (left, pdb 1ssh, (41)), all the peptides are docked to the receptor, converging to the same position. For the failed prediction (right, pdb 1awr (42)), note the peptide segments pointing away from the receptor structure. **(E)** Detection of binding pocket residues using AF2 and PFPD.

#### AF2 predicts the structure of many peptide-protein interactions at high accuracy, comparable to PFPD

Application of AF2 on the *non redundant set* reveals that 25% of the interactions are modeled within 2.0Å and almost 50% are modeled within 5.0Å RMSD (**Figure 1A**, lower panel; unless otherwise noted, RMSD values in the text refer to RMSD calculated over the backbone atoms of peptide interface residues; the best RMSD model among the 5 generated models was selected; for CAPRI assessment values, see **Supplementary Figure S2**). When calculated over all atoms of peptide interface residues, 25% of the interactions are modeled within 3.0Å RMSD. This demonstrates that without any dedicated training or further finetuning, AF2 peptide docking works at high, or at least at medium resolution for a remarkably high fraction of interfaces.

We note that this prediction is possible despite the fact that the MSA does not include any information about peptide sequence conservation (data not shown). This performance is similar to performance on the set of interactions without known motifs, but inferior to modeling of interactions with known motifs, where 50% of the interactions are modeled within an impressive 1.5Å RMSD (on the *motif* and *non-motif PFPD sets*). This suggests that interactions with known motifs can be modeled at higher accuracy. Of note, for the *motif set* performance is comparable to PFPD (where we select the best RMSD model among the top10 cluster centers, as in (21)), while PFPD performs significantly better for interactions with no reported motif (the *non-motif set*) (**Figure 1A**, dotted corresponding lines). Importantly, PFPD and AF2 results show no correlation in performance (**Figure 1B**), indicating that a future combination of the two approaches can significantly boost performance even further.

In contrast to PFPD, using AF2 for peptide docking includes modeling of both the peptide and the receptor. Reassuringly, the performance calculated over the full interface (*i.e*., interface residues of both peptide and receptor) is similar to the one of the peptide, thanks to highly accurate modeling of the individual receptor as well as the peptide structures (**Figure 1C**). (Note, that the low RMSD values for the peptides (**Figure 1C**) can be misleading, since due to their shortness, the maximum difference after superimposing only the peptides is inherently low). **Figure 1D** shows an example of accurate modeling, and another where AF2 fails, easily identified by the poly-glycine linker throwing the peptide segment into space.

We observed that in many cases the models correctly identify the binding pocket, but show errors of peptide rotation or translation (which can result in an RMSD >10Å for a 7-8-mer peptide). These models can still be very useful for the identification of interface residues (**Figure 1E;** see Methods for pocket definition). Interestingly, both AF2 and PFPD recover more of the binding site than one may expect merely by examining the RMSD. At least on this small set, the slightly lower recovery for AF2 is due to finding the right pocket but predicting small translations relative to the native peptide orientation, while PFPD tends to flip the peptide by 180°, still identifying the correct pocket residues. The identification of these interface residues (and hotspots therein) provide a good starting point for further study using low throughput experiments (8).

#### Motif residues are well modeled and can be identified based on the lDDT profile

Given the particularly good performance of AF2 for interactions of proteins with peptides containing a known binding motif (**Figure 1A**, blue lines), we were interested to determine whether we could infer a motif based on our predictions. For this, we inspected performance at the residue level. **Figure 2A** shows heat maps reflecting the per residue RMSD, together with information about motif residues for the *motif PFPD set*. As can be seen, in around 67% of the cases, motif residues show considerably lower RMSD values. Interestingly, for some of the peptides in the *non-motif PFPD set* (**Figure 2B**), we could identify a similar pattern. Reassuringly, we found that the interaction between yeast MAPK Fus3 bound to a peptide derived from MAPKK Ste7 (pdb 2b9h), has a known binding motif (43) that we missed in our previous study (21).

**Figure 2.**
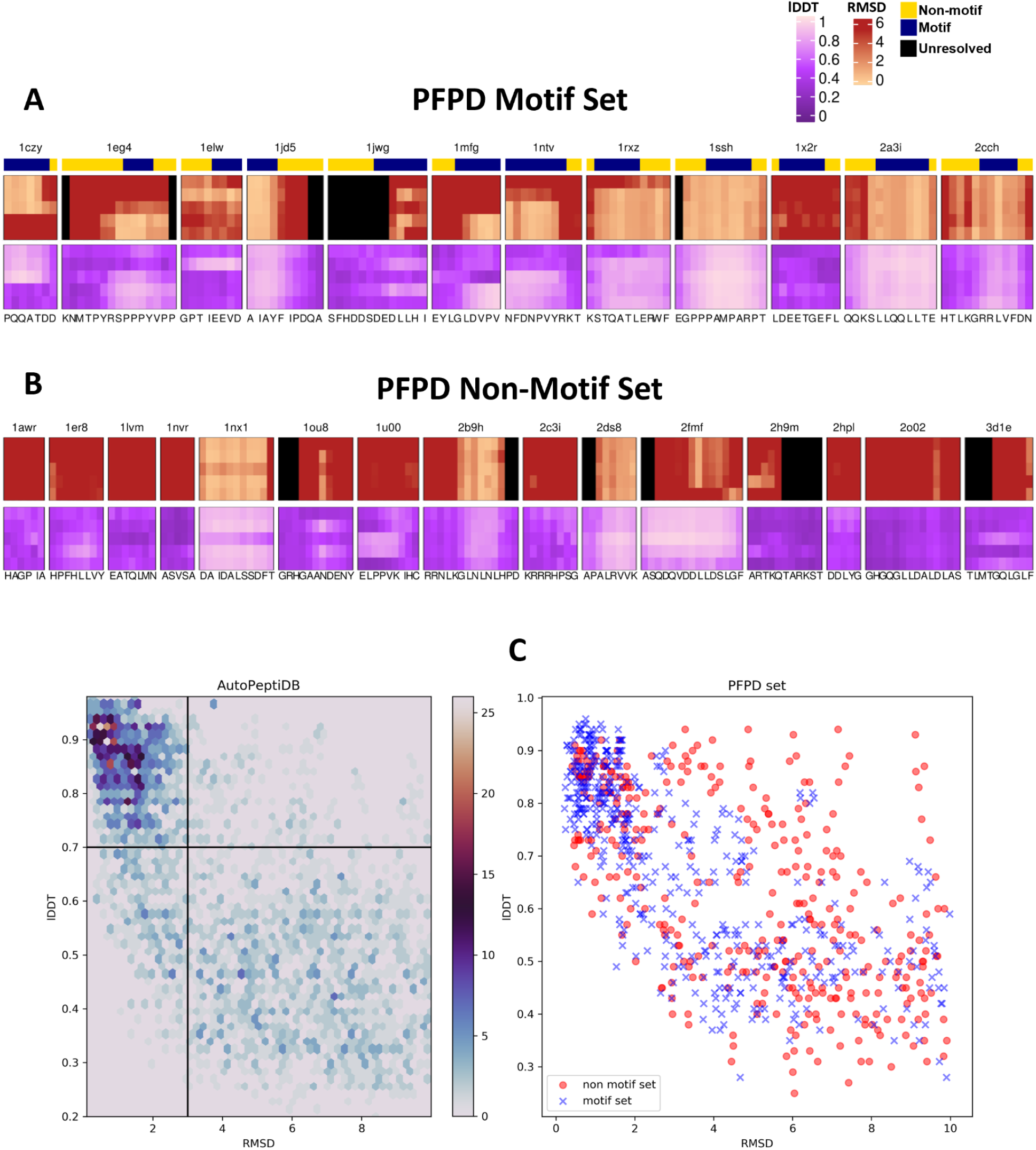
Per-residue RMSD and lDDT heatmaps highlight accurate modeling of the motif region of the peptide by AF2, and suggest a way to identify potential novel peptide binding motifs. **(A)** Heatmaps of per residue RMSD for peptide-protein interactions with known motifs (*motif PFPD set*). The top bar highlights the motif region in blue (compared to the pale yellow colored non-motif residues). The next rows show corresponding lDDT heatmaps. Color scales are indicated to the right (brighter is better in both cases), peptide sequences are annotated at the bottom, each row corresponds to the network parameters used to generate the model (1 through 5). **(B)** corresponding heatmaps for peptide-protein interactions with no reported motif (*non-motif PFPD set*). Note 2b9h, for which reinspection of the literature reveals that a motif was missed in our previous study (and thus, 2b9h should be assigned to the motif set). **(C)** Correlation plots of lDDT *vs*. RMSD values, for the *non-redundant AutoPeptiDB set* (Left), and *motif* and *non-motif PFPD sets* (Right). Plots represent all the per-residue values of the AF2 models (all 5, only residues of the peptides are shown). For the *non-redundant AutoPeptiDB* plot, clusters of points are represented by regional hexagons, colored according to the amount of samples (single residues) in the cluster. In PFPD sets, each dot represents one residue.

Thus, if the correct peptide conformation is available, and therefore, RMSD values can be calculated, novel motifs could be identified based on our structural models. Unfortunately, however, in a real world scenario the peptide structure and the corresponding RMSD values of the models are not known. Luckily, AF2 provides as output also for each model a residue-level confidence estimate, lDDT (Local Distance Difference Test (44)). Inspection of the corresponding heatmaps reveals considerable correlation between the two measures (**Figures 2A,B**). A plot of RMSD and lDDT values for all peptides predicted in this study reveals that this is a general feature: lDDT values above 0.7 consistently represent predictions within 3.0Å RMSD, while values below predominantly reflect predictions above 3.0Å RMSD (**Figure 2C**). This suggests that AF2 predictions may be used to reliably identify correct predictions, and more importantly, previously unidentified motifs. We demonstrate that quite a few longer stretches of amino acids are modeled at high resolution (**Supplementary Figure S4)**, providing a good starting point to look for such new motifs. Further investigation of this avenue is underway.

#### Differences in performance of AF2 and PFPD

In the following we highlight two specific examples in which the two methods show strikingly different performance (highlighted with circles in **Figure 1B**).

##### (1) PFPD outperforms AF2 - 2o02 (14-3-3 protein - exoenzyme S derived peptide)

The receptor 14-3-3 structure is comprised of 9 alpha-helices, with the helical peptide binding within a shallow groove between helices 4 and 9 (45). PFPD positions the peptide in the correct binding site with good RMSD (3.1Å) (**Figure 3A,** left). As for the AF2 model, the peptide is located in the binding site, but flipped and slightly translated: while the nearest residue (Asp427 for 2o02) has an RMSD of 3.4Å, the N-termini are far apart (RMSD ~ 30Å) (**Figure 3A**, right). This is one of several examples in which AF2 finds the site, but positions the peptide only approximately in the binding site (see above).

**Figure 3:**
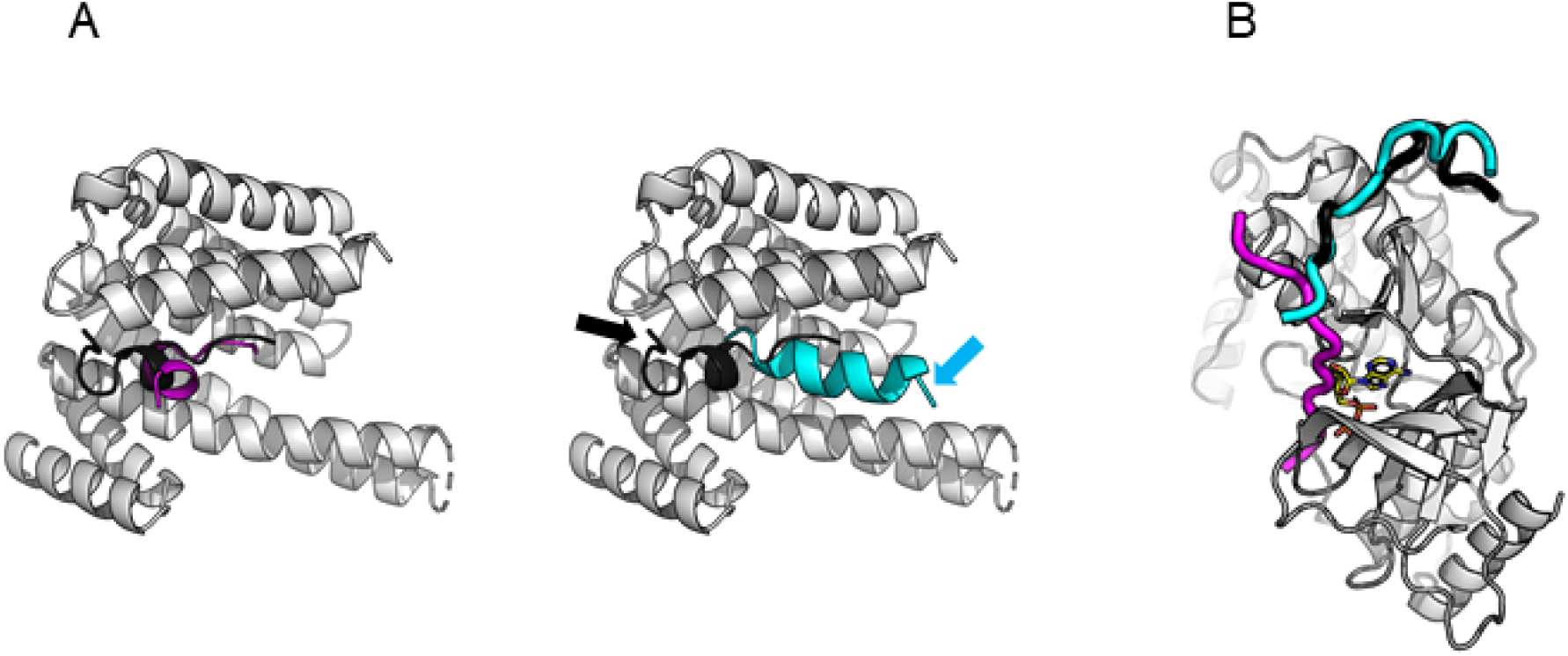
Examples of large variance between AF2 and PFPD predictions. **(A)** PFPD outperforms AF2, interaction between 14-3-3 protein and exozyme S derived peptide (pdb 2o02)The N-termini in the AF2 model and the crystal structure are indicated by arrows, to highlight the direction of the peptide. **(B)** AF2 outperforms PFPD, interaction between MAPK Fus3 and a peptide derived from MAPKK Ste7 (pdb id 2b9h),Peptides in the solved crystal structure are shown in black, the AF2 peptide in cyan and the PFPD peptide in magenta. Only the crystal structure receptor is shown (white) for clarity (the predicted receptor structures are modeled at high accuracy, backbone RMSD=1.2Å).

##### (2) AF2 outperforms PFPD - 2b9h (yeast MAPK Fus3 bound to a peptide derived from MAPKK Ste7)

Here we examine a Mitogen Activated Protein Kinase (MAPK Fus3) binding a peptide derived from its upstream activator MAPK Kinase (MAPKK Ste7) (43). AF2 performance is very good (RMSD=2.4Å), and all models converge to the correct binding site and orientation. In contrast, the corresponding PFPD prediction misses the correct site (RMSD=23Å), positioning the peptide in the ATP binding site. This is a flaw of running PFPD after removing ligands (**Figure 3B**).

#### Modeling peptide binding-induced conformational changes

One of the most challenging tasks in protein docking is to model conformational changes that occur upon binding. Peptide-protein docking approaches have been developed to efficiently and accurately deal with the peptide flexibility, since in most cases no structure is available for the peptide, in contrast to the receptor that adopts a defined conformation in its free, unbound form. However, what happens in cases where the receptor too changes dramatically upon binding? Given the above successful performance of AF2 to predict both receptor and peptide conformation, we hypothesized that these cases would particularly benefit from this new approach where peptide binding is described as part of the folding process. In the following we describe two such examples, demonstrating the ability of AF2 to model peptide binding even upon large changes in the receptor.

##### (1) Estrogen receptor alpha ligand-binding domain bound to peptide derived from nuclear receptor cofactor 2 (residues 686-698)

In this example, the unbound conformation (pdb: 3ert, (46)) positions its c-terminal helix within the peptide binding site in the bound conformation (pdb: 2b1z, (47)). For peptide binding, this c-terminal helix has to move out (by >15Å). We modeled both the bound and the unbound structures, in an attempt to obtain a full picture on the biases and performance. The AF2 model of this peptide-protein complex was spot-on, modeling the correct receptor conformation, and positioning the peptide accurately in the binding site (RMSD < 1.0Å for all models, **Figure 4A**).

**Figure 4:**
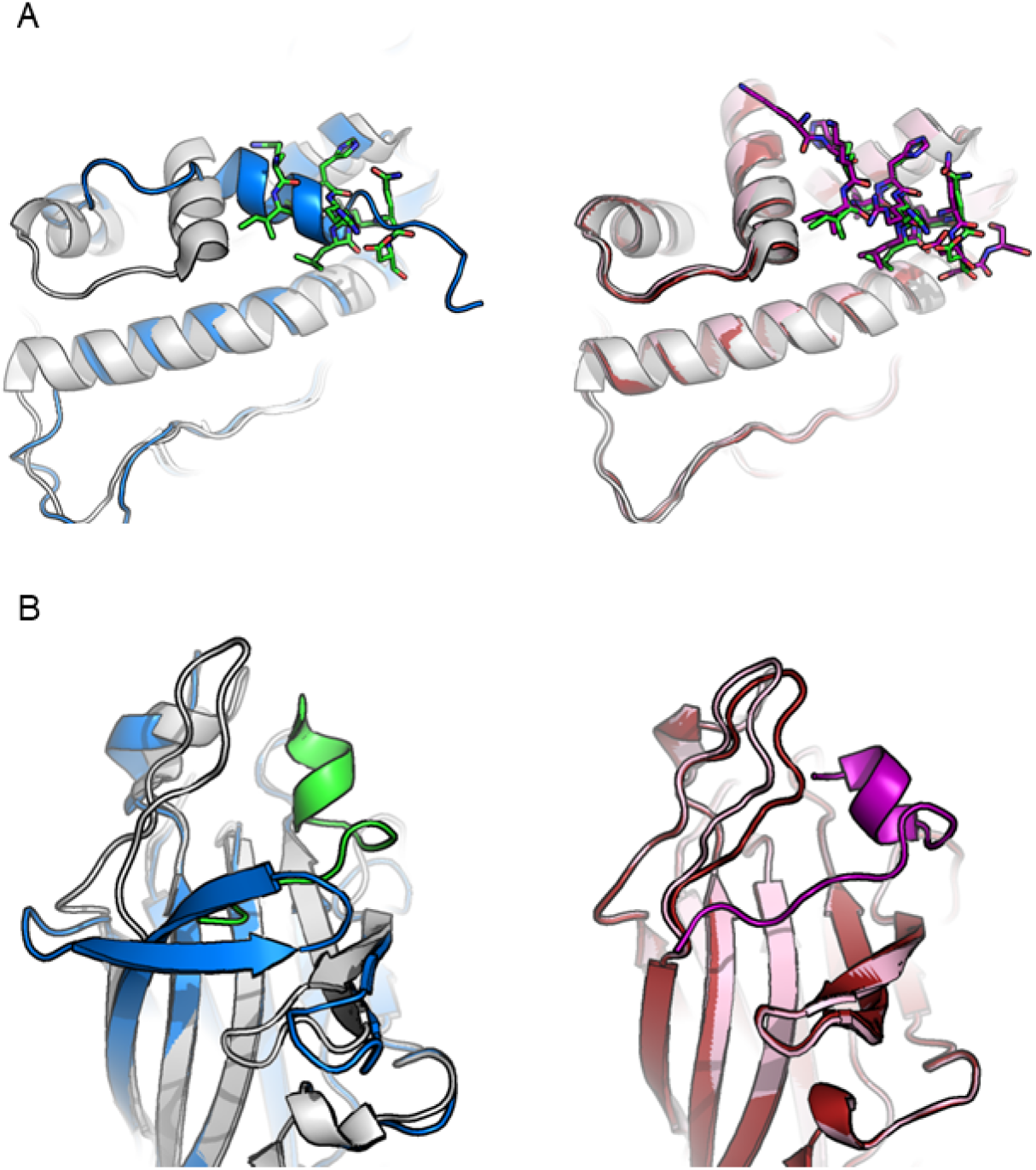
Successful modeling by AF2 of interactions involving significant change in receptor conformation. AF2 models the bound receptor conformation even for the unbound structure, thereby allowing accurate modeling of peptide binding in cases where traditional peptide docking would have failed. **(A)** Estrogen receptor alpha - peptide derived from nuclear receptor cofactor 2. **(B)** EphB4 receptor - ephrin-B2 antagonist. *Left*: Solved crystal structures of the unbound (blue) and bound conformation (receptor in white, peptide in green). *Right*: AF2 model (unbound receptor: maroon, bound receptor: pale pink, peptide: magenta), compared to bound conformation (white). See Text for more details.

This could be due to the fact that our model of the unbound receptor also recovered the *bound* conformation, rather than the published unbound conformation. This indicates that the bound conformation was likely learned, and AF2 predicts the bound conformation by default, even without the presence of the peptide.

##### (2) EphB4 Receptor tyrosine kinases in complex with an ephrin-B2 antagonist peptide

In this similar example, the J–K loop forms a β-hairpin in the unbound conformation (pdb: 3etp, (48)), which upon peptide binding becomes disordered and assumes a flexible loop conformation (pdb: 2bba, (49)). Here again, our unbound model recovers the *bound* structure, with the J–K loop demonstrating a flexible conformation (**Figure 4B**). In this case, only one bound model finds the correct binding position and conformation of the peptide, yet all models place the peptide in the binding site.

To summarize, we show here that AF2 can model not only monomer structures but also many of the interactions between peptides and protein receptors, in particular when a peptide binding motif is available, and even in challenging cases where the monomer changes its conformation upon peptide binding.

## Discussion

In this work we have applied the AF2 protein structure prediction protocol (without any further task-specific optimization) to predict peptide-protein complex structures, at an accuracy comparable to that of state of the art protocols specifically developed for the task of peptide docking. In the following we highlight main insights gained from this study.

### (1) A new efficient approach for peptide-protein docking

High quality predictions were obtained for a large number of protein-peptide complexes (**Figure 1**). AF2 has the advantage of being faster than established protocols such as PFPD - with no trade-off between model quality and runtime. An additional advantage is that AF2 only requires sequences as inputs; no structural information is needed (but can be provided). Finally for AF2 predictions, failures are often easily identified as structures in which the peptide does not interact with the receptor, but rather points out into space.

In turn, among the cases where the peptide is predicted to interact with the receptor, the diversity of interfaces is usually low. While this characteristic is advantageous for reducing false positives, it does not allow for wide sampling of conformational space and assessment of the energy landscape (as is possible with PFPD).

### (2) Peptide binding as monomer structure complementation

The fact that AF2 was trained and tested on monomeric structures, but works for the modeling of peptide-protein interactions, indicates that peptide-receptor binding can indeed be seen as a complementation of the final structure of a monomer.

### (3) Performance is most probably not due to bias in training

As previously mentioned, AF2 was trained on individual chains and thus, as expected, the models of the receptor are indeed very precise (**Figure 1C**). It could however be questioned, whether AF2 successes in the modeling of peptide-receptor complexes could also be due to some “memorization” of interfaces. Our results suggest that this is not the case, since in many cases, while accurately modeling the receptor, and often identifying the correct site, AF2 is not able to generate high resolution models of the bound peptide. Still, AF2 succeeds in peptide docking, indicating that the underlying principles for peptide-protein interactions were well captured and learned - again supporting the view of peptide-protein docking as a protein folding problem.

AF2 was trained on structures collected until 4/2018. In order to further remove possible biases, we also calculated performance on the subset structures released after 8/2018 in the *non-redundant AutoPeptiDB set* (35/162 structures), revealing similar performance (data not shown). In principle however, these cases might well share a homologue in the training set, and thus, this comparison may be biased. Nevertheless, new structures with unrepresented folds bound to peptides are scarce, thus this set would not be fit for a more stringent assessment. As mentioned before however, what counts are the interfaces between peptide and receptor, and AF2 was not trained for that.

To summarize, although remarkable, performance of AF2 is not good enough to assume some hidden overfitting during the training process that we are not aware of.

### (4) Revisiting the role of MSA information in successful modeling

Yet another surprising feature is the ability of AF2 to model peptide-protein complex structures without available MSA information for the peptide. This is particularly surprising since the cornerstone for accurate AF2 predictions is learning residue conservation and co-evolution through contextual processing of MSAs, and benchmarked performance was shown to decrease significantly with decrease in the number of effective alignments for a query (32). The impressive success for peptide docking, albeit completely lacking MSA coverage for the peptide side (in the context of the complex as modeled in this study), is non-trivial. This is yet another indication that the essence of peptide binding can be attributed to implicitly capturing as a substep of folding.

### (5) Modeling large conformational changes upon peptide binding

The advantage of this new protocol lies in its potential to also model considerable conformational changes upon binding. This is due to folding both the receptor and the peptide simultaneously. This would be of special importance in cases where binding induces conformational changes to the receptor (**Figure 4**). This is also an advantage over template-based methods - AF2 can dock peptides to proteins for which close homologues are not available in the PDB (actual performance on this task is yet to be assessed). Nevertheless, it is of course advisable to coerce prior structural constraints on the peptide, receptor or both (and not only use a sequence) to maximize the quality of the prediction.

### (6) Identification of motif instances

Short linear motifs play an important part in binding partner and substrate recognition between proteins. The Eukaryotic Linear Motif resource (ELM, the largest collection of such binding motifs (50)) has more than 3700 annotated instances so far, and this number is continuously growing. Discovering binding motifs and validating them is tedious, although in recent years several high-throughput experimental methods were developed for their detection (51,52).

There are several ways to predict a favorable binding motif for specific interactions or identify new instances of a known motif based on sequence information (53–56). Methods that are based on a solved receptor structure (19) or protein-peptide complex (53,57–60), can be quite effective and provide clues about binding preferences that can later be experimentally validated. However, they mostly require a solved structure of the specific protein and tend to work poorly when modeled based on homolog receptors.

Surprisingly in around ⅔ of the cases, per-residue RMSD, and more importantly for prediction, lDDT values, correctly identify the motif residues within a peptide (**Figure 2**). This is important, since in high-throughput experiments such as phage display (61), a longer stretch of binding peptides is detected, without information about the exact location of the motif within, and often without information about their binding site on the receptor structure. Using AF2 for docking these peptides can be a rapid way to process these results and identify previously unknown motif instances together with their probable binding site and conformation. Especially for peptides without resolved binding motifs, for which AF2 currently performs less well, it might be interesting to investigate whether inclusion of local MSA information extracted from the MSA of the full source protein could improve accuracy of the peptide conformation and the overall accuracy of the complex.

### (7) Fine-tuning will further improve AF2 peptide protein docking

We have presented here an initial, straightforward adjustment of AF2 for peptide docking. Further fine-tuning will without doubt improve the protocol and expose new features that contribute to successful modeling. Parameters to calibrate include the linker length, amino acid composition and its location (current default parameters are 30, poly-G and attachment to the C-terminus of the receptor, respectively). Alternatively, the protocol can be changed to completely remove the linker. Additionally, more sophisticated approaches to MSA generation might result in improved docking and better motif detection. Finally, the orthogonality of performance of AF2 and PFPD (Figure 1B) bears a clear promise for improved peptide-protein docking by combining these approaches.

The experiments reported in this paper and others pave the way towards exciting new avenues for peptide-protein docking and the study of peptide-mediated interactions in general. We believe that by using such approaches, many of the long existing obstacles of the field could be overcome, allowing the study of many more biological systems at high structural resolution.

## Methods

### Generation of a non-redundant dataset of peptide-protein complex structures

For robust assessment of a modeling protocol, it is important to generate a non-biased, non-redundant dataset. For this we have developed the AutoPeptiDB server (39), an automatically updated database of curated peptide-protein interactions. This database is about to be published, but we provide here the details of its generation. For ease of curation and initial analysis, the PDB was queried for entries with 2 chains only, and filtered for those having possible protein-peptide interactions according to the following criteria: **1.** One chain must be over 30 amino acids long, and one chain must be between 4 and 25 AA long. **2.** The peptide chain must have at least 2 residues within 4A distance from the protein chain, yielding a total of 16,931 structures from 1102 ECOD domains (62). Once possible interactions are identified, the following filters are applied: **1.** Remove structures with peptide residues annotated as UNK, **2.** pdb-range and seq-range fields must agree on the indices of the receptor domain according to ECOD annotation (62) **3.** Symmetry operations (from the PDB entry) were applied on the asymmetric unit and checked for possible crystal contacts that may affect the bound conformation of the peptide. Crystal contacts were defined as cases where at least 20% of the peptide residues are in contact with symmetry mates. Finally, the candidates were clustered (within each ECOD domain) according to sequence identity with the threshold set at 30%. Sequence identity was calculated using k-mer distance from the scikit-bio python package and a single representative was selected arbitrarily from the largest cluster of each ECOD domain, yielding a non-redundant set of 162 protein-peptide interactions (complexes that crashed due to their size were omitted).

#### Sequence collection and formatting

The sequences of the receptor and peptide to be modeled were extracted from the SEQRES lines of the PDB files, to account for the expressed construct rather than the structure resolved in the pdb. Unknown residues and terminal modifications were removed. Then, the sequences of each pair were concatenated with a linker of 30 glycine residues, in the order of (N-terminus)receptor-linker-peptide(C-terminus).

#### Structural modelling with AF2

Modelling was performed using the publically available AF2 repository (32), with each of the 5 trained model parameters. The input included the query sequence and MSA from MMseqs2 (63), **without using any templates**. No additional refinement was performed on the models.

Both MSA generation and AF2 predictions were run using publicly available Jupyter notebooks (34) on Google Colab GPUs, slightly modified for batch runs. The modifications did not affect running parameters.

### Analysis of models

#### Global RMSD calculations

Backbone and all-atom RMSDs of the peptide interface residues and the whole interface were calculated using Rosetta FlexPepDock (release 2020.28 (64)). For the models to be comparable via other community-wise used metrics, CAPRI interface metrics (65) were also extracted from the calculated scores (see **Supplementary Figure S2**). Peptide and receptor structure RMSD for backbone atoms and all atoms were calculated using PyMOL python API (v2.2.0), after separately superimposing them onto their native equivalent.

#### Binding pocket calculations

Binding pockets on the receptor were defined as those residues that have at least one backbone atom located within 8.0Å to a peptide backbone atom. The calculations were performed with a PyMOL script (66).

#### By-residue RMSD calculations

Model complexes (protein-peptide) were superimposed onto the native complex as described in the previous paragraph. RMSD was computed using BioPandas python module (67) for each peptide residue pair (model-native), skipping residues that were unresolved in the native structure. Atoms lacking in the models (such as OXT) were also ignored.

#### By-residue lDDT predictions

We extracted the per residue lDDT prediction values from the b-factor column of the structural models output by AF2 (described in (32)).

### Comparison to PIPER-FlexPepDock

Complexes of the motif and non-motif sets were modeled using PFPD as was benchmarked and described previously (21). For each complex, top 10 cluster representatives by FlexPepDock reweighted score were selected for comparison with AF2. Note, that for this assessment, we used the non-redundant set (27 complexes) consisting of two subsets, one with and one without reported motifs (*PFPD motif* and *non-motif sets*), as described therein.

### Visualization

Visualizations were performed with custom R and Python scripts, using packages ComplexHeatmap (68), ggplot2 (69), matplotlib (70) and PupillometryR (71). To visualize structures, we used PyMOL (66).

## Acknowledgements

We are deeply grateful to Sergey Ovchinnikov and Martin Steinegger, and anyone else that has helped provide notebooks to run AF2. This work was supported, in whole or in part, by the Israel Science Foundation, founded by the Israel Academy of Science and Humanities (grant number 717/2017 to O.S.-F.). J.K.V. is supported by Marie Sklodowska-Curie European Training Network Grant #860517. The authors have no competing interests.

## Supplementary Figures

**Supplementary Figure S1.**
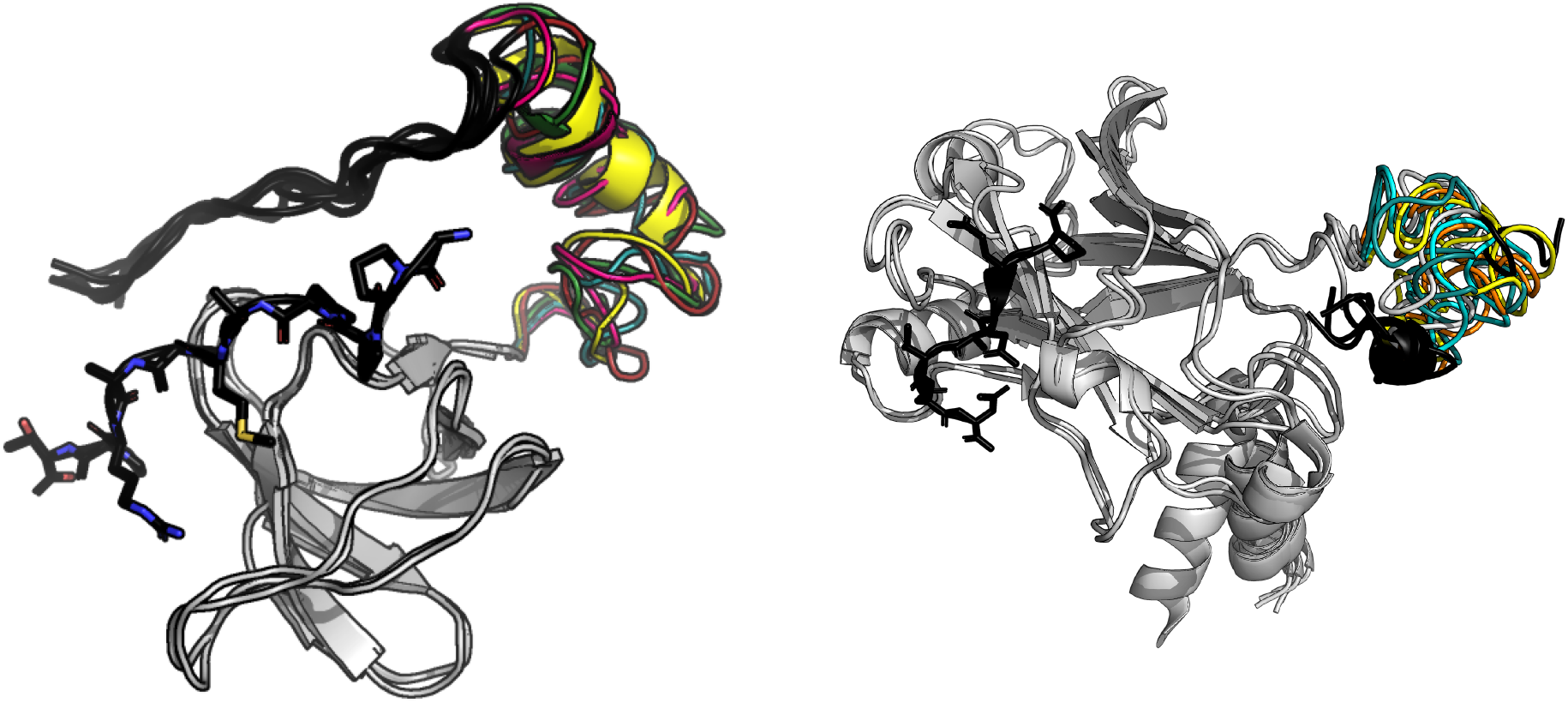
RoseTTAFold attempts to fold the polyglycine linker. Peptide-protein complexes 1ssh (left) and 1czy (right) modeled with RoseTTAFold. The receptor structures are modeled well (light gray, all aligned to the crystal structure), yet the peptides do not reach the binding site (peptides are in black, the peptide from the crystal structure shown in sticks). The polyglycine linker (30 glycines) is folded into an alpha-helix (left) or a highly disordered globule (right), none of which is appropriate for completing the task of peptide docking. AF2 treats polyglycine differently (compare to the AF2 predictions shown in **Figure 1D)**.

**Supplementary Figure S2.**
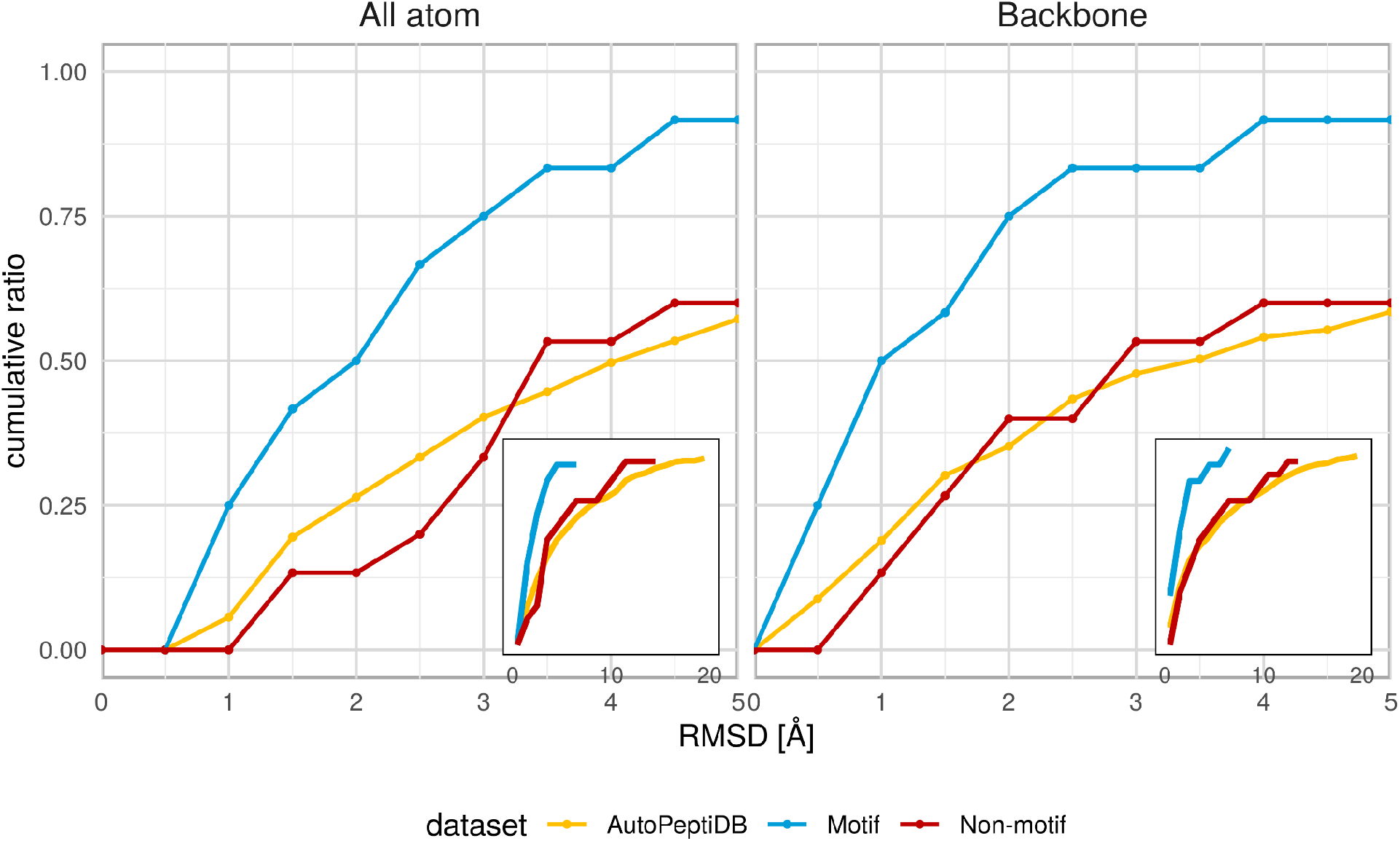
Interface RMSD calculated by CAPRI measures (receptor interface is defined as having at least one heavy atom within 4.0Å distance to a peptide heavy atom and native and model interfaces are superimposed onto each other. Coloring as in **Figure 1.**

**Supplementary Figure S3.**
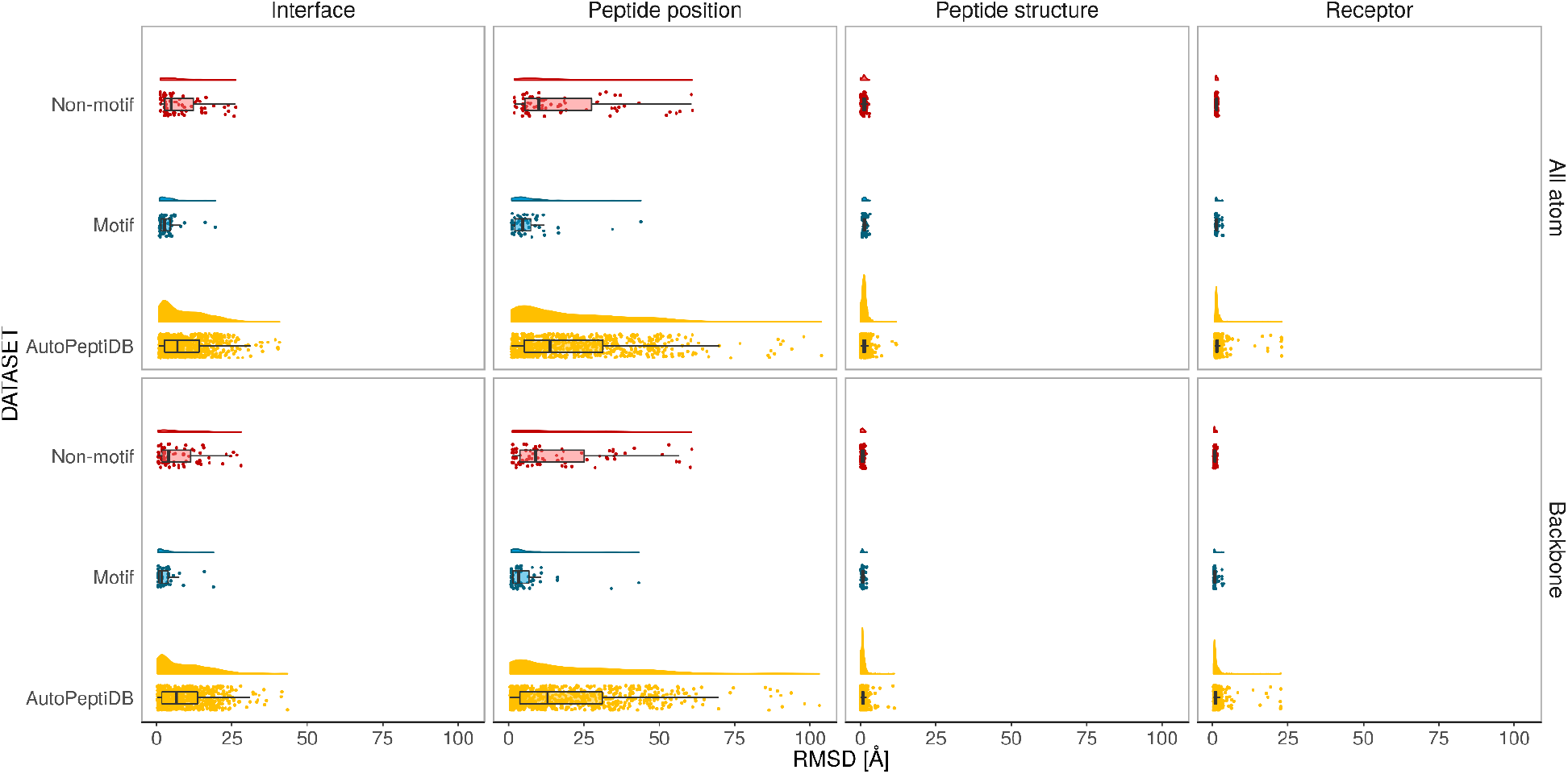
(Accompanies **Figures 1A&C**). Distribution of RMSD values for predicted structures (values >20 were capped to 20). Coloring scheme as in **Figure 1**).

**Supplementary Figure S4.**
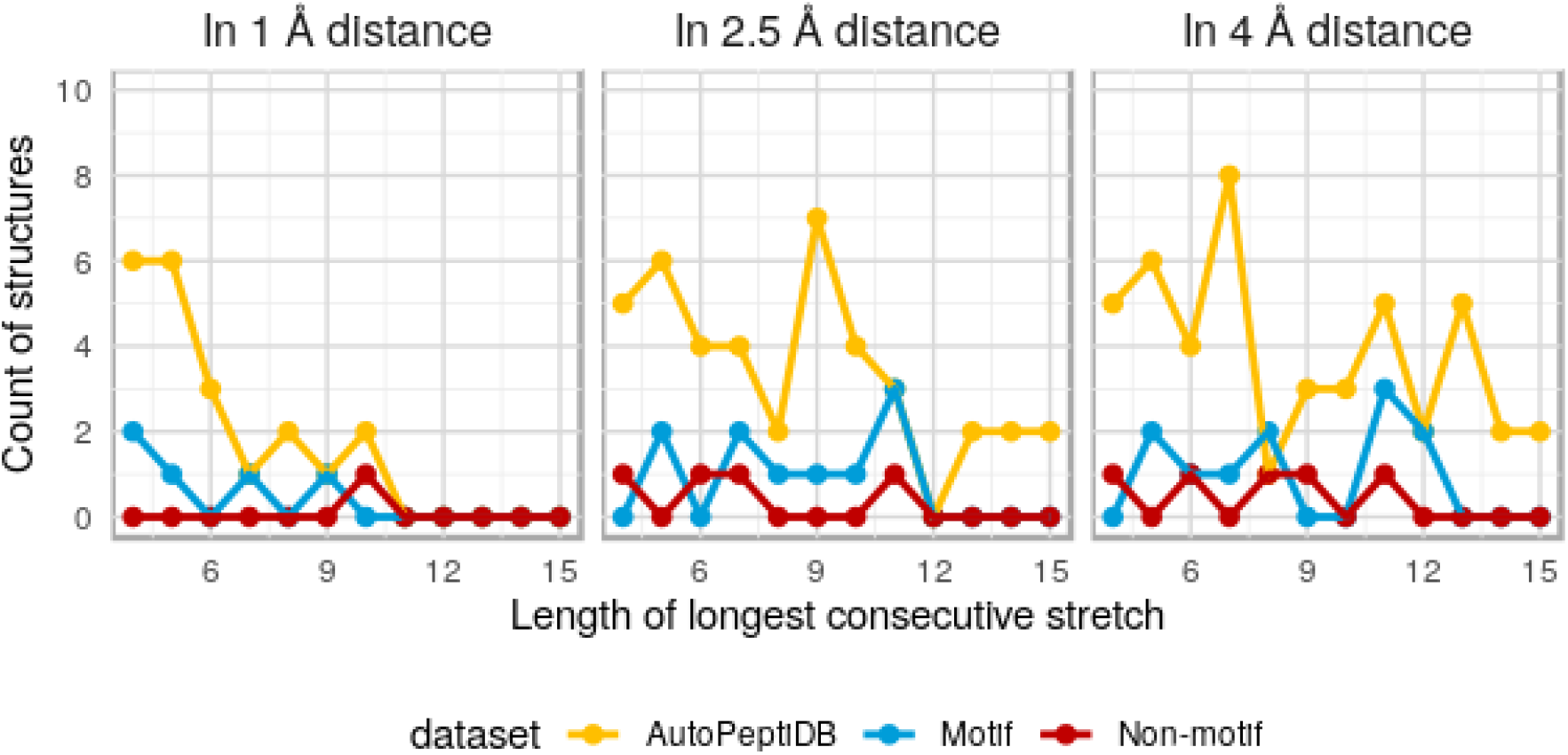
Longest consecutive stretches of well predicted residues. Coloring scheme as in **Figure 1**.

